# Dynamics of pollen tube growth and embryo sac development in Pozna Plava plum cultivar related to fruit set

**DOI:** 10.1101/208108

**Authors:** Milena Đorđević, Radosav Cerović, Sanja Radičević, Dragan Nikolić, Nebojša Milošević, Ivana Glišić, Slađana Marić, Milan Lukić

## Abstract

A newly released, late ripening plum cultivar Pozna Plava sets fruit poorly, although it produces high quality fruit. This study aimed to evaluate which factors in the reproductive process could be related to the lack of fruit set. In two consecutive years, establishment of a suitable polleniser and the stage of ovule development at anthesis as well as initial and final fruit set have been studied. In addition to this, the impact made by temperature fluctuations on the interaction between male gametophytes and female sporophytes was also analysed. Growth of the pollen tubes in the style and penetration into the nucellus as well as fruit set were more effective in cross-pollination than in open and self-pollination. A relative delay in ovule development was observed, and most ovules had an embryo sac with eight nuclei. Considering the results of the quantitiative parameter study of pollen tube growth in the ovary as well as the results of the stage of ovule development, a conclusion can be made that this cultivar is characterised by an extremely short effective period of pollination.

## 1. Introduction

In order to enable realisation of the full yield potential of a cultivar, it is necessary to understand various aspects of fertilisation biology (Cerović and Mićić 1996). The progamic phase, which begins when a pollen grain reaches the stigma, is characterised by specific interactions between the male gametophyte and the female sporophyte (de Graaf et al. 2001). Following the adhesion of pollen on the stigma, the female sporophyte takes over the subsequent processes, including hydration, germination, growth through the conducting tissue of the style and guidance to the micropyle, until the final interaction with the female gametophyte (Palanivelu and Tsukamoto 2011). Based on the analysis of pollen tube growth in sour and sweet cherry, Stösser and Anvari (1981) reported that pollen tubes reach the base of the style in two days, whereas it takes them six to eight days to reach the ovules. Also, Cerović (1994) showed that early penetration of pollen tubes in sour cherry ovules occurred on the fourth day after pollination. In sweet cherry, a faster growth of pollen tubes in the style than in the ovary was detected, suggesting a possible inter-dependence between the number and speed of pollen tube growth on one side and the sporophytes of the mothers genotype on the other (Radičević et al. 2016).

The maturity stage of an ovule is considered an important factor in the process of fertilisation. An immature ovule during the full blooming of sour cherry (Furukawa and Bukovac 1989) and apricot (Alburquerque et al. 2002) was reported as a possible cause of poor fruit set. Normal and mature embryo sacs at the time of full bloom is thought to be directly associated with the success of fertilisation of sour cherry (Cerović and Mićić 1999) and apricot (Alburquerque et al. 2002). Another aspect of fertilization biology is the EPP concept, which was introduced by Williams (1970) in order to assess the difference between the duration of vitality of ovules and the time needed for the pollen tubes to reach the ovules. In general, the EPP was demonstrated as a good framework for the identification of factors that limit the fruit set. Sanzol and Herrero (2001) pointed out that EPP is conditioned by three main processes: stigmatic receptivity, pollen tube kinetics and ovule development, which are under the influence of a number of factors, such as temperature, pollen quality and chemical treatments. A poor fruit set in some plum cultivars may occur due to genetic predisposition resulting in the development of irregular embryo sacs, as well as low temperatures during the blooming period. This may lead to weak growth of the pollen tubes (Thompson and Liu 1973). Apart from these, a poor fruit set may occur as a result of a reduced EPP (Williams 1970; Stösser and Anvari 1982). A short interval of stigma receptiveness is certainly a disadvantage, since it may result in a low fruit set.

One of the characteristics of global climatic warming is the increase in the frequency of extreme temperatures, which have an impact on the reduced success rate of fertilisation, thereby causing a low fruit set (Tubiello et al. 2007). High temperatures occurring at the time of full bloom have a bigger impact on the female gametophyte, causing a rapid degeneration of the embryo sac and ovule, compared to the impact made on the germination of pollen at the stigma and the growth of pollen tubes in the style (Beppu et al. 2001). Conversely, low temperatures occurring at the time of bloom tend to reduce the pollen tube growth rates in the ovary, thereby shortening the effective pollination period (Sanzol and Herrero 2001). Hall (1992) highlighted that the reproductive phase is when plants are most sensitive to temperature fluctuations.

The present paper was undertaken primarily to discover the nature of irregular cropping of a newly released, late-ripening and high-quality plum cultivar Pozna Plava in agroecological condition of Western Serbia. In order to investigate the reproductive characteristics of Pozna Plava, establishment of a suitable polleniser, the effect of ovule maturity at anthesis on the fruit set and ovule development were assessed.

## 2. Materials and Methods

### 2.1. Plant material

Pozna Plava is a newly realised cultivar within a plum breeding programme at Fruit Research Institut, Čačak (FRI), Republic of Serbia, which has been derived from self-pollination of Čačanska Najbolja (Lukić et al. 2016; Đorđević et al. 2016). This cultivar is protected in the EU area under the name Čačak Späthe, in colaboration with nursery Ganter OHG Markenbaumschule, Wyhl, Federal Republic of Germany. The study of reproductive properties of Pozna Plava was conducted in three pollination variants: self-, cross- and open pollination. In a cross-pollination variant, three economically important pollen donor cultivars, i.e. Čačanska Najbolja (developed at FRI, Čačak, Republic of Serbia), Hanita and Presenta (both developed at University of Hohenheim, Federal Republic of Germany) were used. The experiment was carried out over a two-year period at Ljubić facility of the FRI (located in southwestern Serbia, 43°N latitutde, 20°W longitude, and 224 m altitude). The orchard was established in 2002, with cultivars grafted on a myrobalan (*Prunus cerasifera* L.) rootstock, with a spacing of 6.0 × 5.0 m and an open vase training system.

### 2.2. Pollen viability *in vitro*

To study pollen viability *in vitro*, branches with flower buds from four plum cultivars were sampled in the field, at late balloon stage, during two consecutive years (2010–2011). The anthers were collected and placed on paper at room temperature until dehiscence. The pollen was used for testing pollen viabily *in vitro* and for pollination of isolated flowers of Pozna Plava. The viability tests were then immediately performed. Pollen was scattered in Petri dishes on germination medium consisting of 1% agar and 12% sucrose, and kept at 25°C. A pollen grain was considered as germinated according to method described by Galleta (1983). Counts of germinated pollen grains were made under a light microscope Carl Zaiss (Jena, Federal Republic of Germany), while they were photographed using the Olympus BX61 microscope (Tokyo, Japan). For each cultivar, three replicates per plate with approximately 100 pollen grains were counted. From these data, the percentage of germinated pollen grains of each genotype were estimated.

### 2.3. Pistil sampling

The study of pollen tube growth in the Pozna Plava pistils was carried out in all three pollination variants. In field conditions, flowers on all selected trees of Pozna Plava were emasculated and isolated by bagging shortly prior to anthesis. Pollination of emasculated flowers in self- and cross-pollination variants was performed at the beginning of full flowering. Again, these branches were isolated with bags, which were then removed 10–15 days after pollination. Forty pistils in all pollination variants were sampled at 2 day intervals, starting from the time of full bloom, until 10 days after full bloom (DAF). To fix the samples, pistils were immediately soaked in FPA solution (70% ethanol, propionic acid and formaldehyde, 90:5:5 percentages by volume) and stored at 4°C. Aniline blue staining was used for this study (Preil 1970; Kho and Baër 1971). Using a razor blade on a microscope slide, the style was cut at the base and removed from the ovary, before being opened along the suture and covered with the cover slip. The ovary was opened and, to enable better observation of pollen tube penetration in the micropyle and nucleus, the ovule was cut longitudinally–tangentially. The numbers of pollen tubes in the upper part of the style and ovary were counted on the fluorescent microscope (Olympus BX61) under ultraviolet light, using Multiple Image Analysis in AnalySIS software. The quantitative efficiency of the pollen tube growth in the style and ovary for each variant of pollination was examined by determining the average number of pollen tubes and the dynamics of their growth in certain regions of the pistils. Counting of pollen tubes in the style was performed in the upper third section, while in the ovary, it was conducted in the region of placenta. In all the pollination variants, the number of pollen tubes from five fixing intervals is shown in the form of a mean value. The dynamics of pollen tube growth is shown by the percentage of pistils with the penetration of the longest pollen tube in certain regions.

For examining the ovule development at anthesis, flowers from open pollination and emasculated unpollinated flowers were used during 2008 and 2010. Every other day, starting from the onset of full bloom until 10 days later, 20 flowers of both variants were picked randomly. Pistils were fixed in FPA and stored at 4°C. Ovaries were dehydrated in an ethyl alcohol series and then embedded in paraffin wax. Serial sections of 10 μm were mounted on slides impregnated with an adhesive of gelatine. Samples were stained with safranin, crystal violet and green light SF, as described by Gerlach (1969) and sealed with a cover slip using canada balsam as an adhesive. Observation and photography was performed under an Olympus BX61. Primary ovules were studied and the developmental stage was established. According to Dyś (1984) and Furukawa and Bukovac (1989), in both variants, the following ovule developmental stages were established: embryo sac with two nuclei; embryo sac with four nuclei; embryo sac with eight nuclei; embryo sac with eight nuclei but non-fused polar nuclei; embryo sac with eight organised nuclei and fused polar nuclei. Embryo sacs with degenerated antipodes at five-nuclear stages were also counted as functional. The ovule developmental stage in both variants was expressed as the percentage of ovules studied in the sample.

### 2.4. Fruit set

To determine the fruit set in all pollination variants at the time of full bloom, 3–4 branches were marked with approximately 100 flowers per branch. Initial fruit set was determined 4–5 weeks after the pollination and final fruit set was determined at the beginning of ripening.

### 2.5. Examination of air temperature

In order to record the climatic conditions in 2010 and 2011 during the full flowering phase, daily temperatures were collected. Full flowering time was established according to Wertheim (1996), i.e. as the date when 80% of flowers were opened.

### 2.6. Statistical analysis

The data were statistically analysed using Fishers two-factor model of variance analysis (Fischer 1951). Arcsin square-root data transformations were performed. Regarding the factor for which the F test showed statistical significance, an individual test was performed using LSD test at *P ≤ 0.05*. Pearson correlation and determination coefficients were used to determine mutual dependence between certain tested parameters. Fixed nonlinear regression equations were used to determine the dependence of normal embryo sacs on the respective study days. Statistical analyses were performed using SPSS statistical software package, Version 8.0 for Windows (SPSS. Inc., Chicago, IL).

## 3. Results

### 3.1. Pollen viability *in vitro*

Establishing the germination potential of pollen in different fruit types is an important characteristic reflecting pollen quality, considering that efficient fertilisation requires a high percentage of pollen germination activity. The weakest pollen germination in both years of the study was determined in Pozna Plava, reaching an average of 17.3%. In the other pollenisers used in the experiment, the average germination rate was between 42.3%(Čačanska Najbolja) and 46.5% (Presenta) (Fig. 1).

**Fig. 1.**
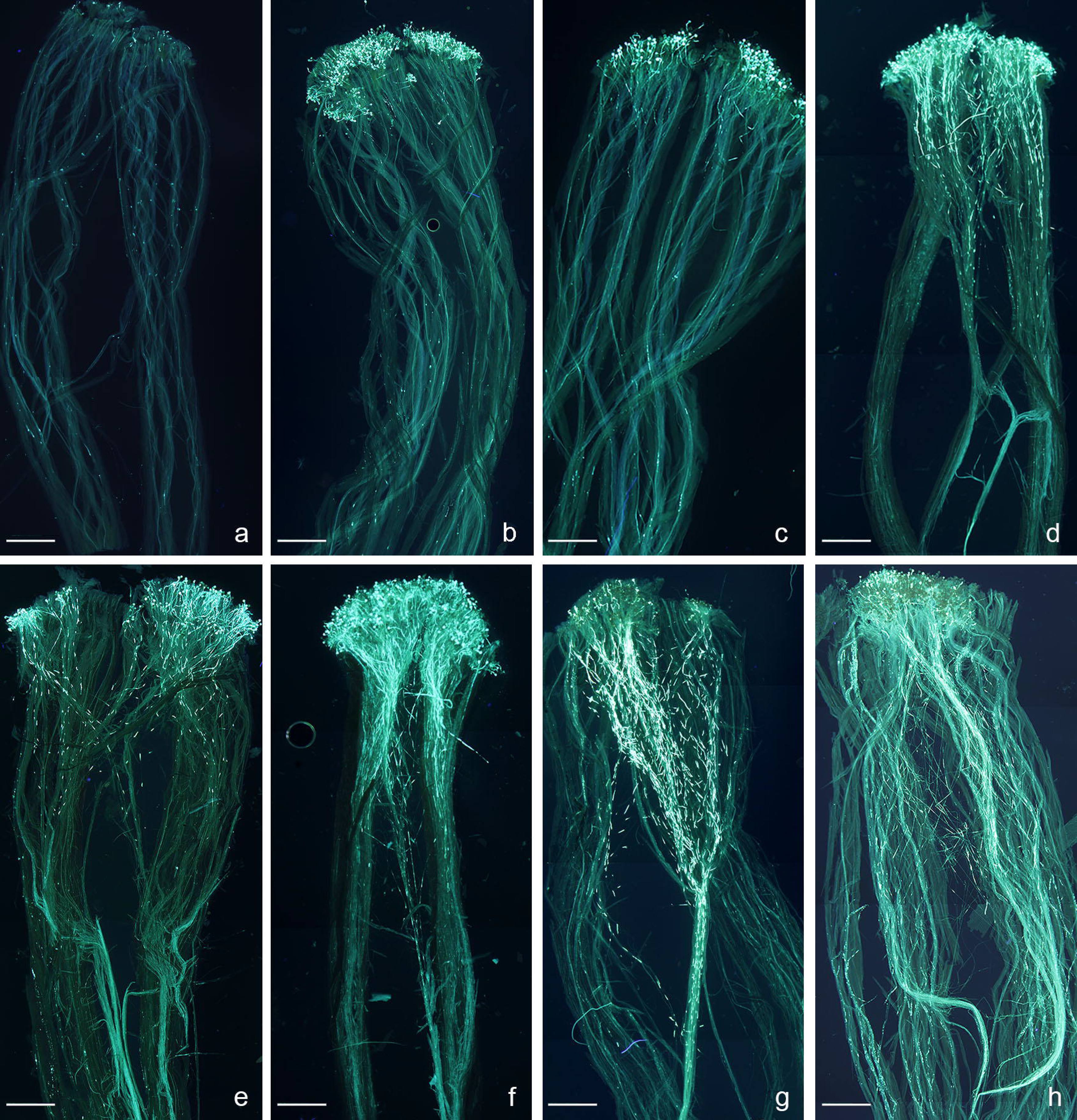
Pollen germination *in vitro*.

## 3.2. Average number of pollen tubes in the pistils

The highest average number of pollen tubes in the upper third of the style (61.15) and the ovary (3.15) of Pozna Plava was observed with Hanita as polleniser (Table 1), whereas the lowest number of pollen tubes was found in the open pollination variant (17.94 and 1.33).

**Table 1.**
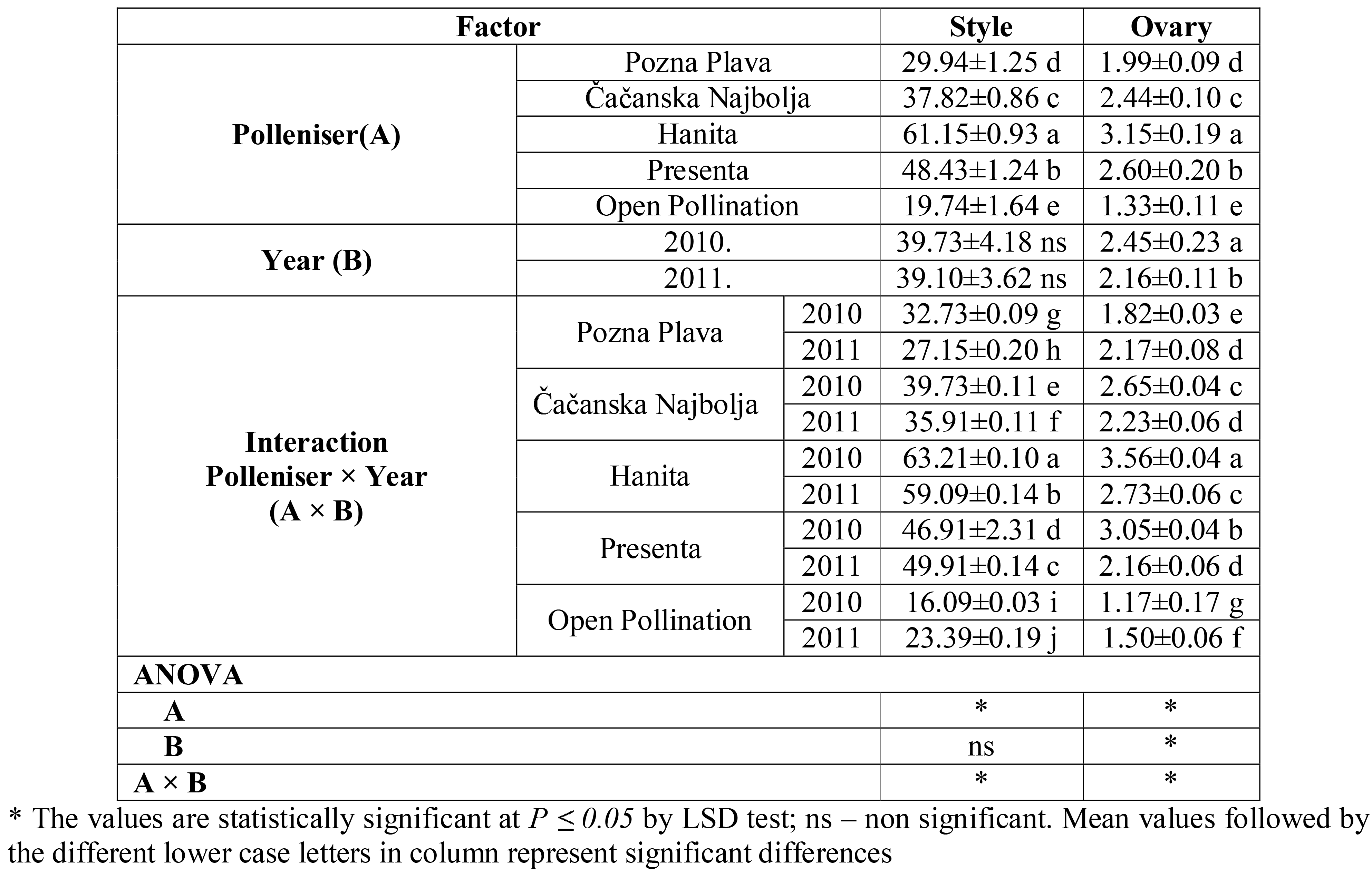
Average number of pollen tubes in certain regions of the pistils in the Pozna Plava.

## 3.3. Dynamics of growth of pollen tubes in the ovary

During the two-year period, the longest growth of the pollen tubes was obtained in the self-pollination variant, whereas the weakest growth was recorded in the open-pollination variant (Fig. 2). In the first year of the study, the longest growth of the pollen tubes occurred within the first 10 DAF, with Hanita as the polliniser (48.17% of the pistils with the penetration of the pollen tubes into nucellus). The weakest growth of pollen tubes in all the days of the trial during 2010 was observed in the open pollination variant (Fig. 3 a-h and Fig. 4 a-c).

**Fig. 2.**
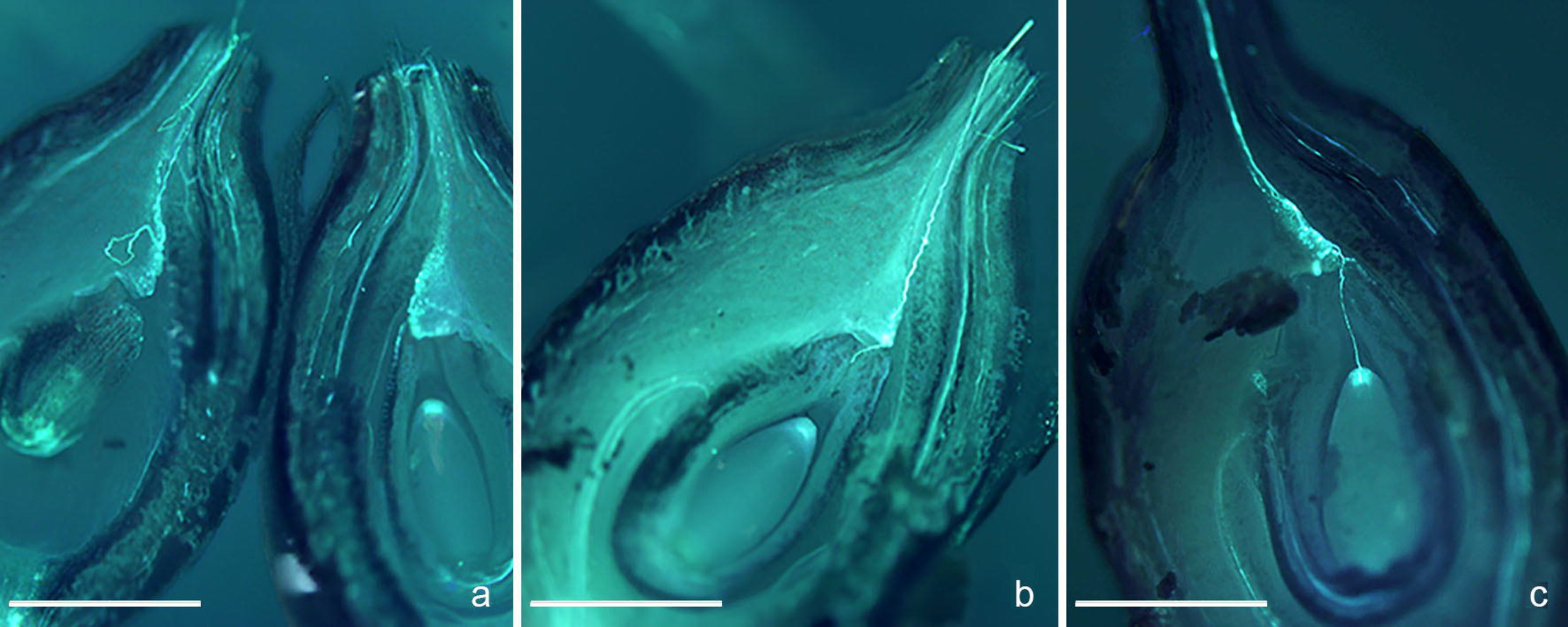
Dynamics of pollen tubes growth in certain regions of the Pozna Plava pistils, in the examined pollination variants, during the two-year period (*Stu*, *Stm*, *Stb* – upper, middle and lower third of the style; *O* – ovary tissue; *M* – micropyle; *N* – nucellus).

**Fig. 3.**
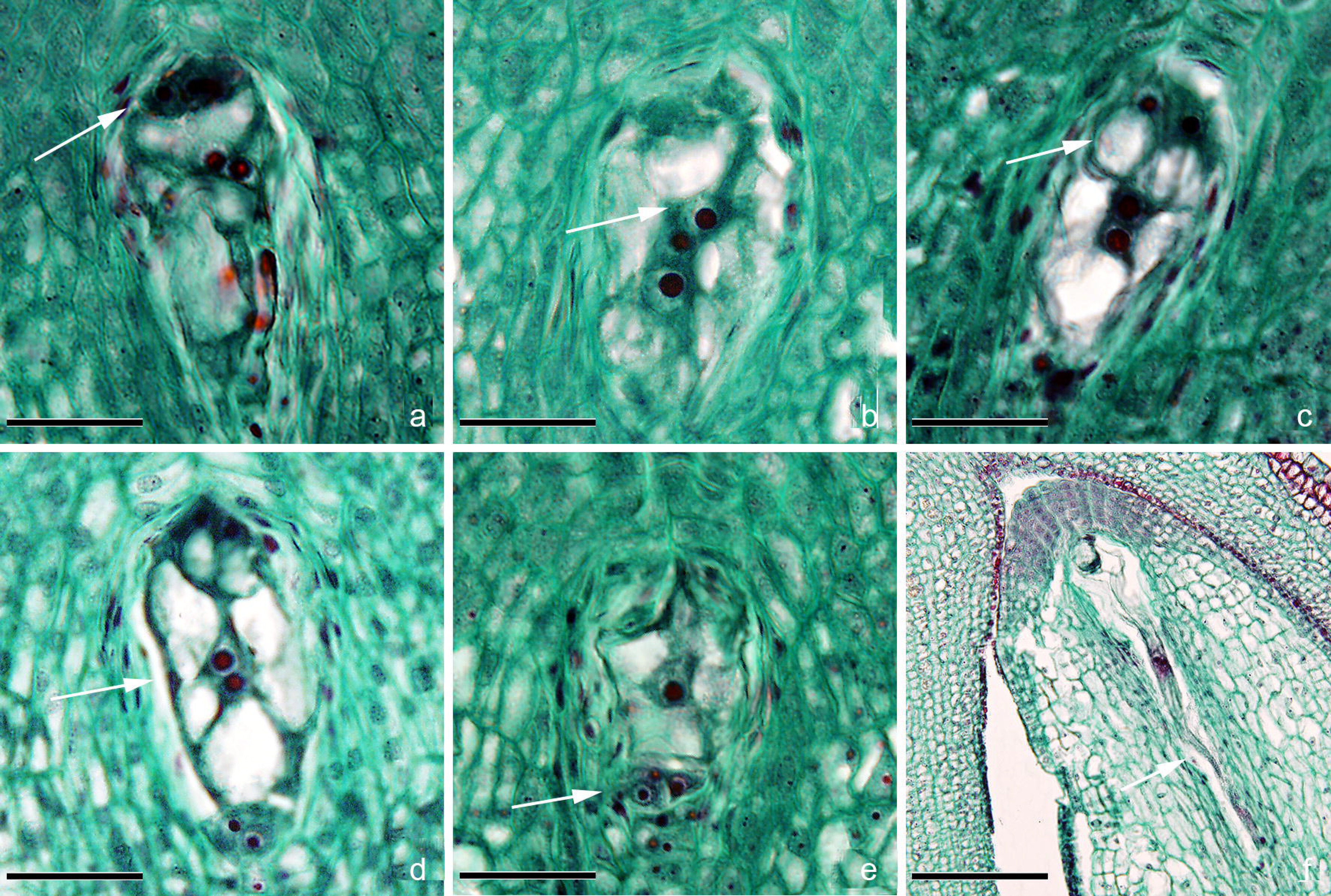
Growth of pollen tubes, in the analysed pollination variants, in the style of the Pozna Plava: a-b) on the 2 DAF; c-e) on the 4 DAF; f) on the 6 DAF; g) on the 8 DAF; h) on the 10 DAF Scale bars=1mm.

**Fig. 4.**
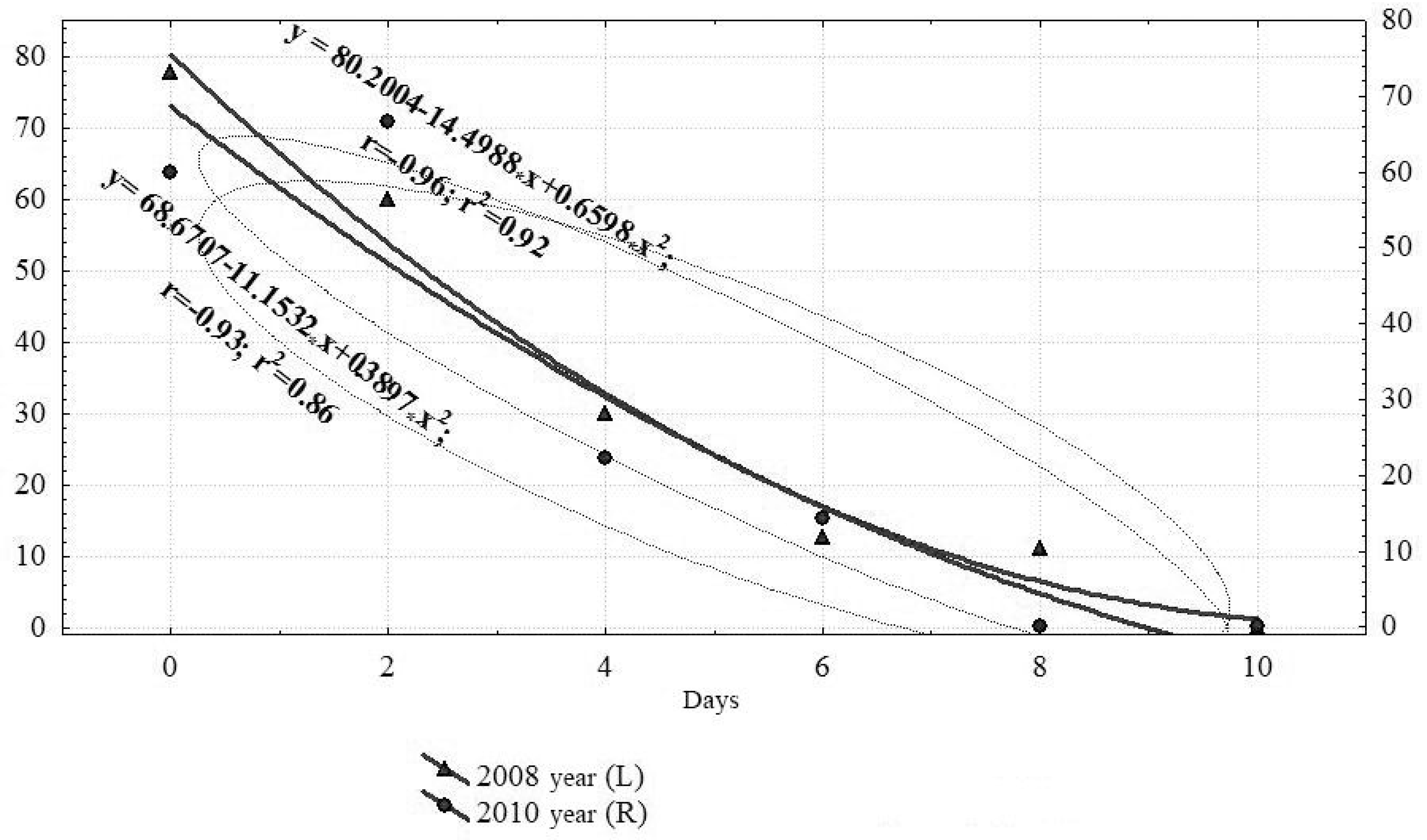
Penetration of pollen tubes in certain regions of the Pozna Plava ovary: a) obturator (self-pollination variant); b) micropyle (pollination with Hanita); c) nucellus (pollination with Presenta) Scale bars=1mm

The weakest growth of pollen tubes in the first few days of fixing was determined in the open or the self-pollination variant, whereas on 10 DAF, it was weakest in the open pollination variant.

The value of 0.76 of the Pearson correlation coefficient indicates a high positive correlation between the number of pollen tubes in the upper third of the style and the number penetrating the ovary. A high positive correlation (0.72) was also established between the number of pollen tubes in the ovary and the number of the same tubes penetrating the ovule nucleus on the tenth day after pollination (Table 2).

**Table 2.**
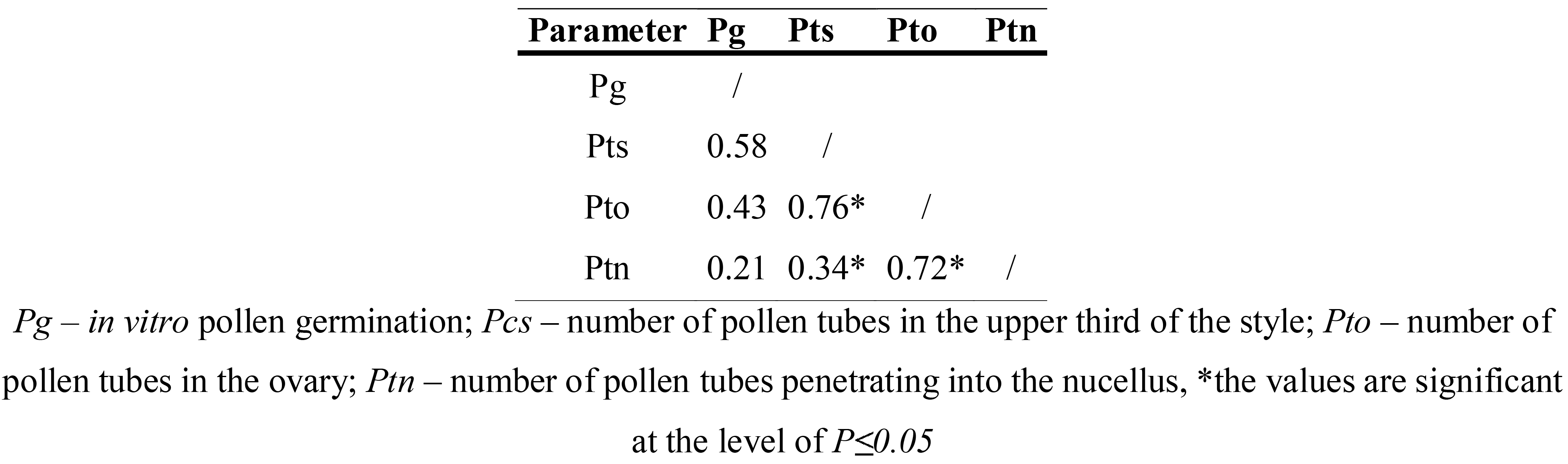
Pearsons coefficient of correlation between germination rate polena *in vitro*, number of pollen tubes in the ovary and penetration of pollen tubes in ovule

## 3.4. Histological study of the ovary

During the two experimental years, the obtained results always revealed two ovules, which – given minor variations – were of approximately the same size at the onset of the full flowering period. The larger and better developed ovule was taken as the primary, and the other was considered as the secondary ovule.

Formation of an embryo sac in Pozna Plava occurred in accordance with *Polygonum* type. There were four clearly distinctive types of cells in the embryo sac: the egg cell, the synergid cell, the central cell and the antipodes (Fig. 5a). At the time the embryo sac became mature, the cytoplasm and the nucleus was located in the chalazal part, crescent-shaped, while the remaining part was filled with a large vacuole (Fig. 5b).

**Fig. 5.** The functional stages of embryo sac and its elements at longitudinal views: a) early 8–nuclear stage; b) appearance of egg cell; c) pear-like form of synergids; d) appearance of central cell; e) antipods; f) elongation of the embryo sac Scale bars=0.1 mm

The synergids were located in the micropillar segment of the embryo sac, and the application of triplet colouring demonstrated that their cytoplasm was thickest and mostly concentrated around the nucleus, with the remaining part towards the chalazae section occupied by a large vacuole (Fig. 5c). The central cell was the largest cell in the embryo sac, with 80% of its contents filled with vacuoles at the time of maturity (Fig. 5d). On average, 3–4 large vacuoles can be found in the central cell. Two polar nuclei were located in the central part, closer to the micropyle, while the larger part of the cytoplasm was located in the peripheral section, although it was possible to see the cytoplasm threads in the middle section. Three antipodes were located in the chalazae section of the embryo sac. They were ephemeral and somewhat larger than the other cells surrounding them (Fig. 5e). The antipodes could only be seen in the first few days of full flowering.

Analysis of embryo sacs in both variants during a two-year period, in the full flowering phase, has shown that these were in the early phase of forming a mature embryo sac. In 2008, in the open pollination vaiant, on 8 DAF the proportion of embryo sacs with five nuclei was 12.50%, compared to 2010, when on the same day, the number of embryo sacs in the four-nuclear phase was 25% (Table 3). In both years, on 10 DAF, it was possible to observe gradual elongation of embryo sacs towards the chalazal part (Fig. 5f).

**Table 3.** Normal cytological structure of the embryo in the primary ovule of Pozna Plava flowers in the open pollination variant, in the first 10 DAF in both study years

| Stage | Number of primary ovules with normal embryo sac (%) |  |  |  |  |  |  |  |  |  |  |  |
| --- | --- | --- | --- | --- | --- | --- | --- | --- | --- | --- | --- | --- |
|  | Full bloom |  | 2 DAF |  | 4 DAF |  | 6 DAF |  | 8 DAF |  | 10 DAF |  |
|  | 2008 | 2010 | 2008 | 2010 | 2008 | 2010 | 2008 | 2010 | 2008 | 2010 | 2008 | 2010 |
| 4-nuclei |  |  |  | 10.00 |  |  |  |  |  |  |  |  |
| 8-nuclei | 14.29 |  |  | 10.00 |  |  |  |  |  |  |  |  |
| 8-nuclei embryo sac, early phase | 28.58 | 30.00 | 10.00 | 10.00 |  |  |  |  |  |  |  |  |
| Mature embryo sac | 14.29 | 20.00 | 10.00 | 40.00 |  | 10.00 |  |  |  |  |  |  |
| 5-nuclei embryo sac | 14.29 |  | 50.00 | 10.00 | 60.00 | 40.00 |  | 20.00 | 12.50 |  |  |  |
| 4-nuclei embryo sac |  |  |  |  |  |  | 10.00 |  |  | 25.00 |  |  |
| 3-nuclei embryo sac |  |  |  |  | 10.00 | 10.00 | 50.00 | 20.00 | 50.00 | 37.50 | 50.00 | 20.00 |
| 2-nuclei embryo sac |  |  |  |  |  |  |  | 10.00 | 12.50 | 12.50 | 10.00 | 10.00 |
| Early embryo stage |  |  |  |  |  |  |  |  |  |  |  | 10.00 |
| Number of analyzed ovules | 7 | 10 | 10 | 10 | 10 | 10 | 10 | 10 | 8 | 8 | 10 | 10 |

In the unpollinated flowers higher percentage of embryo sacs in the early phase of forming a mature embryo sac was established in 2008 (Table 4). In 2008, on 2 DAF, the functionality of embryo sacs changed in favour of five nuclei (due to degeneration of antipodes), whereas on the same day in 2010, it was possible to detect early eight-nuclei, eight-nucei and five-nuclei stages of the embryo sac development.

**Table 4.** Normal cytological structure of the embryo in the primary ovule of non-pollinated flowers of Pozna Plava in the first 10 DAF in both study years.

| Stage | Number of primary ovules with normal embryo sac (%) |  |  |  |  |  |  |  |  |  |  |  |
| --- | --- | --- | --- | --- | --- | --- | --- | --- | --- | --- | --- | --- |
|  | Full bloom |  | 2 DAF |  | 4 DAF |  | 6 DAF |  | 8 DAF |  | 10 DAF |  |
|  | 2008 | 2010 | 2008 | 2010 | 2008 | 2010 | 2008 | 2010 | 2008 | 2010 | 2008 | 2010 |
| 4-nuclei | 11.11 |  |  |  |  |  |  |  |  |  |  |  |
| 8-nuclei | 11.11 |  |  |  |  |  |  |  |  |  |  |  |
| 8-nuclei embryo sac, early phase | 55.56 | 50.00 |  | 11.11 |  |  |  |  |  |  |  |  |
| Mature embryo sac |  |  |  | 22.22 |  |  |  |  |  |  |  |  |
| 5-nuclei embryo sac | 11.11 | 10.00 | 60.00 | 33.33 | 30.00 | 22.22 | 12.50 | 14.29 | 11.11 |  |  |  |
| Number of analyzed ovules | 9 | 10 | 10 | 9 | 10 | 9 | 8 | 7 | 9 | 9 | 10 | 7 |

In 2008, there was a drastic drop in the functional five-nuclei stage of an embryo sac, established on the fourth day of the full flowering phase (30.00%). In both years of the study, a smaller percentage of the five-nuclear stage was established on the sixth day, whereas only 11.11% of the samples were found to be in the five-nuclear stage on 8 DAF. In the same period in 2008 and 2010, the embryo sacs in the remaining samples were characterised as having lost their functionality due to cytological changes.

While the established functional stages were presented in a summarised form, the functional dependence on the day of study is statistically best presented in the form of a regression curve (Fig. 6). The obtained values indicate the presence of a high negative correlation between the number of days after the full bloom and the examined functional cytological stage of an embryo sac. The obtained values of the correlation coefficient and determination in 2008 were −0.96 and 0.92, while these values were slightly lower in 2010, reaching the range between −0.93 and 0.86.

**Fig. 6.** Non-linear regression between the functional phase of an embryo sac and the number of days after the full bloom of the non-pollinated flowers of Pozna Plava in the two-year period

Based on the obtained results, the existence of functional stages of the embryo sacs can be assumed to persist until the eighth day after the onset of the full flowering phase.

## 3.5 Fruit set

In both years of the experiment, the highest fruit set of Pozna Plava was obtained in the crosspollination variant, whereas the lowest number of fruit was determined in both self- and open pollination variants. In all the analysed variants, a better fruit set was recorded in 2010 (Fig. 7).

**Fig. 7.** Fruit set of Pozna Plava in different pollination variants

The highest values of the final fruit set in both years were obtained in the cross-pollination variant, followed by the self-pollination variant. The lowest fruit set of Pozna Plava was recorded in the open pollination variant (less than 5%). The obtained values of the Pearson correlation coefficient indicated the presence of a statistically significant impact of the number of pollen tubes penetrating into the nucellus on 10 DAF on the number of initial and final fruit set (Table 5).

**Table 5.** Pearsons coefficient of correlation between penetration of pollen tubes in nucellus 10 DAF, initial and final fruit set

| Parameter | Ptn | Ini | Fin |
| --- | --- | --- | --- |
| Ptn | / |  |  |
| Ini | 0.77* | / |  |
| Fin | 0.44* | 0.39* | / |

*Ptn* – number of pollen tubes penetrating into the nucellus 10 DAF; *Ini* – initial fruit set;
*Fin* – final fruit set; \*the values are significant at the level of $P \leq 0.05$

The correlation between the penetration of the pollen tubes in the ovule and the initial set was relatively high, at 0.77, whereas the coefficient of determination was 0.59. The relationship between the penetration of pollen tubes in the ovule and the final fruit set, as well as the relationship between the initial and final fruit showed inter-connection values (0.44 and 0.39), whereas the determination coefficient for these analysed values was 0.20 and 0.18.

## 3.5. Air temperature

2010, at the start of full bloom and 1 DAF, mean daily temperature was in the approximate range of 11°C. Within the following six days, mean daily temperature ranged between 14.45°C and 17.67°C, before they declined once again to around 11°C.

Higher temperature fluctuations were recorded in 2011. Following the temperature values of 10°C and 16°C at the start of the full bloom, for the following four days the mean day temperature values were in the range between 4.03°C and 8.2°C, before resuming higher values in the range between 11.35°C and 18.17°C. The average mean daily temperature in the period under consideration during 2010 was 14.03°C, compared to 10.63°C in 2011.

## 4. Discussion

### 4.1. *In vitro* pollen germination and part played by the progamic phase on fruit set

In order to attain high and regular yield, understanding of the functional capacity of pollen is highly significant, considering that the *in vitro* behaviour of pollen may be used as a reliable indicator of its behaviour *in vivo*. The pollen germination rate obtained for the plum cultivars assessed in this paper is in accordance with results reported for a large number of plum cultivars and hybrids in the agro-ecological conditions of western Serbia, which were in the range between 25.6% and 45.4% (Đorđević et al. 2012; Glišić et al. 2012). In comparison to our result (44.82%) Milošević (2013) reported a slightly lower value for the *in vitro* pollen germination of Hanita (36.60%).

No correlation was found between the *in vitro* pollen germination rate and the number of pollen tubes in the upper third of the style. Similar results were obtained by Hormaza and Herrero (1996), who reported a better *in vitro* pollen germination rate in the examined sweet cherry cultivars, compared to the initial *in vivo* germination rate. Keulemans (1994) concluded that the *in vivo* pollen germination rate is affected by the temperature, often resulting in lower *in vivo* pollen germination rate values compared to the *in vitro* germination rates of pollen. The decrease in pollen tube penetration in the ovary was between 10 and 19 times lower in the tested pollination variants. The literature on the growth of pollen tubes in the ovary of different fruit species has so far indicated a reduced number of pollen tubes present in the lower sections of the style or in the ovary, such as in apricot (Herrero 1992) and plum (Nikolić and Milatović 2010). Following the analysis of the average number of pollen tubes in the style and the ovary, the largest values were obtained in the cross-pollination variant, while the lowest values were determined in the self-pollination and open pollination variants, in the two years of the study. The obtained results clearly indicated an impact of the polleniser genotype on the tested parameter. With the possible exception of the presence of a certain amount of its own pollen, the reason for the low values of pollen tube average number in the open pollination variant may be low quantitative presence of pollen at the stigma.

Based on the obtained results, it can be concluded that the best growth dynamics were obtained in the cross-pollination variant. A faster growth of pollen tubes in cross-pollination variants was reported in, sour cherry (Cerović 1994), sweet cherry (Radičević 2016) and plum (Jia et al. 2008). Concerning representatives of the *Prunus* genus, the growth of pollen tubes in the style is faster than their growth rate in the ovary, despite shorter spatial distance covered by the pollen tubes in the ovary, compared to the distance in the style (Herrero 1992). In apricot, Rodrigo and Herrero (2002) reported in the pollination variants under consideration that pollen tubes became visible in the style base after three to four days, whereas their penetration in the nucellus could be noticed on the seventh day after pollination. Similar results were obtained in peach, where pollen tubes reached the base of the style within seven days, which was followed by a period of inactivity, lasting until the cells of the obturator entered into the secretion phase (Herrero and Arbeloa 1989).

Based on the two-year study of the ovary in Pozna Plava, including the pollinated and non-pollinated variants, it can be concluded that this cultivar is characterised by a normal morphological structure of the ovary, featuring two ovules in the ovary attached to the placenta wall with the funiculus. The normal eight-nuclear cytological structure of the embryo sac in the open pollination variant was observed in the first few days after full flowering, at the same time as the five-nuclear stage in both years of the experiment. As early as the second day after the start of full flowering, observations revealed a larger number of ovules with the five-nuclear stage of the embryo sacs. In 2008 and 2010, starting from the fourth day after full flowering, an observation revealed the presence of ovules characterised by degeneration of one synergid and a partial or full degeneration of the second synergid. In the literature, the process of double fertilisation is commonly related to degeneration of one of the synergids and the merging of the polar nuclei into a central nucleus. However, in this study a different tendency was observed; a larger number of ovules contained degeneration of both synergids, while a low number of ovules were recorded as having fused the polar nuclei into the central nucleus. According to Yang et al. (2010), degeneration of synergids can be a physical, as well as a physiological process. During 2010, on the tenth day after the onset of full flowering, an observation revealed an early stage of embryogenesis in one of the samples, with the presence of nuclei of the future endosperm.

### 4.2. Fruit set

The weather conditions at the time of pollination impacted the receptiveness of the stigma, fertility of the ovule, duration of its vitality and the fruit set (Furukawa and Bukovac 1989). In addition, there is a large number of genotype-conditioned factors related to flowering biology that also have the potential to impact the fruit set, including, among others, the production and shedding of flower buds, time of flowering, stage of ovule development at flowering time and the germination capacity of pollen (Ruiz and Egea 2008). Higher values of the initial and final fruit set, apart from the open pollination variant, were obtained in the other pollination variants in 2010 compared to 2011. A possible cause for the weaker fruit set in 2011 could be low temperatures occurring at the time of full flowering, which resulted in the slower growth of the pollen tubes.

### 4.3. Temperature fluctuations during the full bloom sub-phase

Air temperature represents a major ecological factor with a direct impact on the pollination and fertilisation process (Irenaeus and Mitra 2013), thereby making a direct impact on the fruit set (Vasilakakis and Porlingis 1985). The impact of high temperatures immediately prior to or during flowering had a negative effect on the progamic phase in the peach, resulting in a weak fruit set (Nava et al. 2009).

A possible explanation for the established differences in the average number of pollen tubes in the upper third of the style ought to be sought in the action of ecological factors, primarily the temperature during flowering phenophase. During 2011, the temperature in the flowering period was extremely low, with a possible decisive impact on the lower average number of pollen tubes in the first three of the pollenisers listed above. The higher average number of pollen tubes in 2011 in the cross-pollination variant with Presenta may be explained by this cultivar originating from a colder climate. At the same time, the same tendency in the open pollination variant can be explained by the lower temperatures in the period of full bloom also causing a prolonged period of flowering in certain plum cultivars, thereby causing their flowering time to overlap with the Pozna Plava flowering time, enabling the availability of larger quantities of pollen. The impact of low temperatures on the dynamics of pollen tube growth in 2011 was not significantly pronouced, since the dynamics were almost levelled. This can be explained by the adaptive capacity of Pozna Plava and Čačanska Najbolja, both domestic by origin, whereas the other two cultivars, Hanita and Presenta originated from a colder climate and showed their adaptavity to low temperatures. This is in accordance with the findings of Hedhly (2005), who pointed out that the pollen–temperature interaction may reflect the origin of the pollen, as well as its adaptation to the prevailing ecological conditions. The low rate of penetration of pollen tubes in the ovule on the tenth day of pollination in 2011 could be attributed to the high temperature fluctuations on the sixth day, where the difference in the daily temperature means reached up to 8°C.

The impact of temperature on various structures or processes during the progamic phase was observed, such as the receptiveness of the stigma (Hedhly et al. 2003), germination rate of the pollen grain at the stigma, the growth of pollen tubes in the style and ovary (Hedhly et al. 2008) and duration of vitality of the ovules (Cerović et al. 2000).

## 5. Conclusion

This study reports a reproductive survey of Pozna Plava in order to solve irregular cropping, which was revealed in this excellent quality plum cultivar.

The results have shown that although Pozna Plava is a self-pollinating cultivar, far better results were obtained in the cross-pollination variant. A markedly poor fruit set in the open pollination variant has revealed the absence of an adequate polleniser in the orchard with the analysed cultivar.

Considering the study results of the embryo sac functional stage of this plum cutivar, as well as the results of the quantitative parameters of pollen tube growth in the ovary, a conclusion can be made that this cultivar is characterised by an extremely short EPP. This highlights a need for a proper choice of two to three pollenisers, which – in addition to the overlap in the full flowering phase with this cultivar – will be able to yield pollen grains of good quality.

Based on the obtained values of the tested parameters of the pollen tube growth in the ovaries of Pozna Plava, the cultivars Hanita, Presenta and Čačanska Najbolja can be recommended as good pollenisers for this plum cultivar analysed in the agro-ecological conditions of Čačak.

## Acknowledgements

This work was conducted under Research Project TRD□31064: Development and preservation of genetic potential of temperate zone fruits, supported by the Ministry of Education, Science and Technological Development of the Republic of Serbia.

